# Age-related changes in noise correlations in a decision-making task

**DOI:** 10.64898/2026.06.27.734969

**Authors:** Fenying Zang, Anne E. Urai

## Abstract

Age-related declines in sensory and cognitive performance may arise from changes in neural variability, including how variability is shared across neural populations. Here, we examined age-related changes in noise correlations, which quantify shared trial-to-trial variability between pairs of neurons, in mice performing a visual decision-making task. We analysed large-scale extracellular recordings from 149 mice (male and female) spanning 3 to 20 months of age, comprising 348,937 simultaneously recorded neuron pairs within 17 cortical and subcortical brain regions. Across the dataset, noise correlations showed expected relationships with firing rate, anatomical distance, and signal correlations, consistent with canonical observations of noise correlations. Age associations with noise correlations varied across regions and were bidirectional: the age-related increase in primary visual cortex replicated previous findings in this region, whereas decreases were observed elsewhere. Evidence for age-related differences in stimulus-induced changes was limited. Together, these findings indicate that ageing is associated with regionally heterogeneous changes in shared neural variability rather than a brain-wide shift.

## Introduction

Age-related declines in sensory and cognitive performance may arise from increased neural variability. Across species, ageing is associated with slower and more variable perceptual decisions across a range of tasks (MacDonald et al., 2006; Li & Rieckmann, 2014; Zang et al., 2026), raising the possibility that behavioural decline reflects a loss of precision in the underlying neural computations. Theoretical and behavioural studies have suggested that age-related increases in neural variability may reduce the effective signal-to-noise ratio within the central nervous system, thereby impairing information processing and cognitive performance (Crossman, E. R. & Szafran, J., 1956; Welford, 1981; Cremer & Zeef, 1987). We previously found that ageing was associated with higher post-stimulus, but not pre-stimulus, single-neuron variability (measured by Fano factor), together with attenuated stimulus-induced variability quenching during visual decision-making (Zang et al., 2026). These findings suggest that ageing may not only increase neural variability, but also alter how variability changes during task-relevant sensory processing.

Beyond variability in individual neurons, neural variability is increasingly studied as structure in how activity co-varies across neural populations. Recent advances in large-scale electrophysiology and imaging have expanded the ability to characterize correlated population activity and examine its consequences for neural coding, computation, and behaviour (Urai et al., 2022; Stringer & Pachitariu, 2024). This focus builds on foundational studies showing that shared variability can shape population coding in ways that are not captured by single-neuron measures alone (Averbeck et al., 2006; Cohen & Kohn, 2011).

Noise correlations, often referred to as spike-count correlations in the literature (*r_sc_*; here termed *r_noise_* to distinguish them from signal correlations), quantify the extent to which trial-to-trial fluctuations in firing rate are shared between pairs of neurons under repeated stimulus presentations (Cohen & Kohn, 2011; Zohary et al., 1994). They matter because shared variability can influence how much information neural populations encode and how effectively that information can support behaviour. Theoretical work further shows that the consequences of shared variability depend not simply on its magnitude, but on its structure: shared fluctuations aligned with the stimulus-coding direction can fundamentally limit the information available to downstream decoders (Averbeck et al., 2006; Moreno-Bote et al., 2014).

Importantly, noise correlations are not static, but are dynamically modulated by cognitive and brain state, including attention, arousal, and task engagement (Cohen & Maunsell, 2009; Ecker et al., 2014; Rabinowitz et al., 2015; Doiron et al., 2016). These changes are behaviourally relevant: experimental studies in behaving animals have shown that improved perceptual performance is often accompanied by reduced noise correlations, including under attention manipulations in the visual cortex (Cohen & Maunsell, 2009) and following perceptual learning (Gu et al., 2011; Ni et al., 2018). Moreover, pharmacological modulation that reduces noise correlations in the visual cortex has been reported to improve visual performance in rhesus macaques (Ni et al., 2022). Together, these findings suggest that shared variability reflects moment-to-moment changes in the coordination of population activity rather than a fixed property of a circuit. From this perspective, ageing can be viewed as a longer-timescale shift in brain and behavioural state, extending the same logic from moment-to-moment fluctuations in attention or arousal to changes that unfold across the lifespan (Urai, 2026). This raises the possibility that shared variability may also be reshaped by ageing.

Existing animal studies suggest that ageing may be associated with increased noise correlations, with evidence reported in both primates and rodents and across visual and auditory cortex (Liu et al., 2025; Shilling-Scrivo et al., 2021, 2022; Wang et al., 2019), potentially reflecting inhibitory circuit decline or altered excitation-inhibition balance. At the same time, the available evidence remains relatively narrow in scope. Most ageing studies have been conducted during spontaneous or passive conditions, with only few experiments measuring noise correlations during active behaviour. Existing studies have also focused largely on specific, mostly sensory cortical, brain areas. Moreover, few of these findings have been replicated across independent laboratories or large standardized datasets, a broader challenge in neuroscience (Botvinik-Nezer et al., 2020). It therefore remains unclear whether age-related changes in noise correlations are robust across datasets and laboratories, and whether they generalize across brain regions during active decision-making.

Here, we leveraged large-scale Neuropixels recordings from younger and older mice performing a standardized visual decision-making task (International Brain Laboratory et al., 2021) to characterize age-related differences in noise correlations during behaviour. Across 348,937 recorded neuron pairs from 17 cortical and subcortical regions, we asked whether age effects were shared across the brain or region-specific, and whether ageing altered stimulus-related modulation of noise correlations. Age associations with pre- and post-stimulus noise correlations were region-specific and bidirectional: the age-related increase in primary visual cortex replicated previous findings in this region, whereas decreases were observed elsewhere. Evidence for age-related differences in stimulus-induced changes was more limited. Together, these findings indicate that ageing is associated with regionally heterogeneous differences in shared neural variability rather than a brain-wide increase.

## Results

We analysed publicly available large-scale Neuropixels recordings from the International Brain Laboratory dataset (International Brain Laboratory, Meshulam, et al., 2025; Zang et al., 2026), obtained from mice performing a standardized visual decision-making task (International Brain Laboratory et al., 2021). Mice were trained to decide whether a visual stimulus appeared on the left or right side of the screen by turning a small steering wheel (Figure 1a). Stimulus contrast varied across trials, thereby modulating task difficulty. The dataset covered a broad continuous age range from 3–20months (Figure 1b), roughly corresponding to a human age range of 23–70 years (Cottam et al., 2025). For visualization only, mice were divided into young and old groups using the dataset mean age of 7.6 months as the cutoff: young mice (N = 97, mean age = 5.50 months) and old mice (N = 52, mean age = 11.56 months). All statistical analyses treated age as a continuous variable.

**Figure 1.**
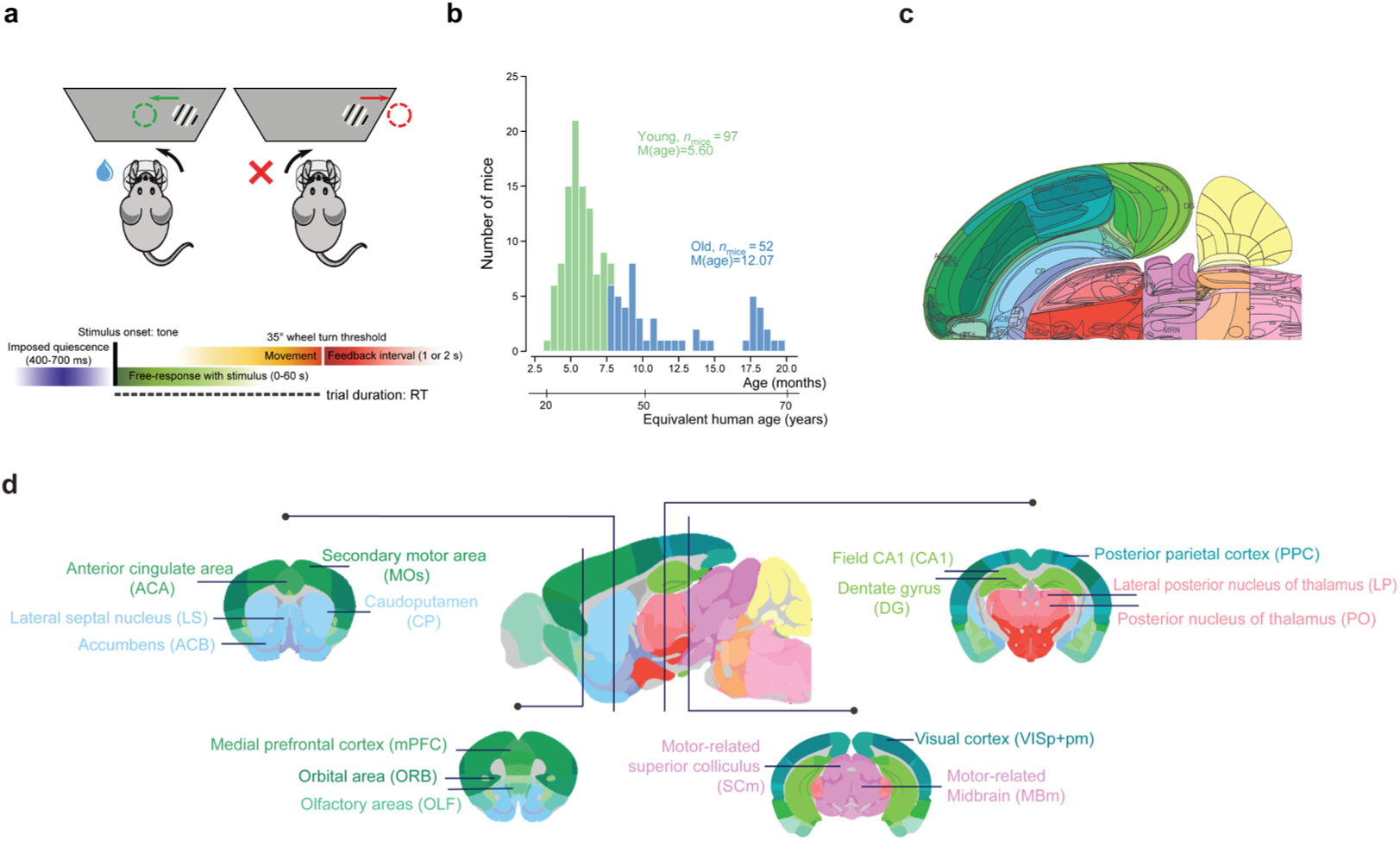
The IBL behavioural task and the datasets. **(a)** Schematic of the task, showing a correct response (the visual stimulus was brought to the centre of the screen) vs. a wrong response (the visual stimulus was moved off the screen). Adapted from [International Brain Laboratory et al.](https://doi.org/10.1038/s41586-025-09235-0); © The Author(s) 2025; licensed under [CC BY 4.0](https://creativecommons.org/licenses/by/4.0/)). **(b)** Distribution of mouse age on the day of neural recording (n = 149 mice; young n = 97; old n = 52; age cutoff: 7.6 months). Group splits were only used for visualization; all statistics used age as a continuous variable. The age range roughly corresponds to a human age range of 23–70 years (Cottam et al., 2025). **(c)** The 2D Swanson flatmap representation of the mouse brain. Labelled regions indicate regions of interest (ROIs), as detailed in (d) and Table 1. **(d)** A 2D-sagittal mouse brain slice and four corresponding coronal slices, adapted from the Allen Mouse Brain Atlas and Allen Reference Atlas – Mouse Brain (mouse.brain-map.org and atlas.brain-map.org), with ROI acronyms and annotations added.

**Table 1.**
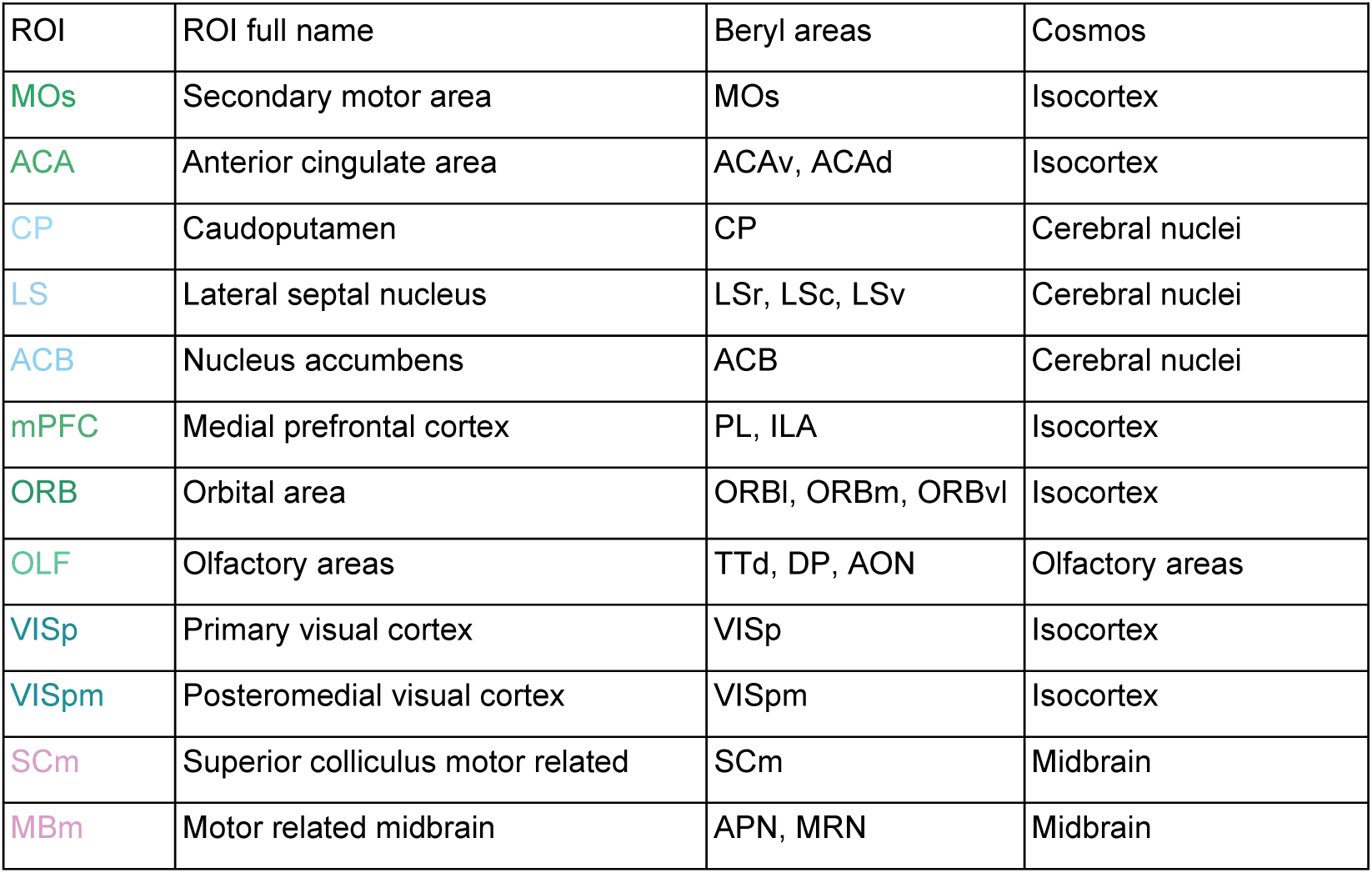
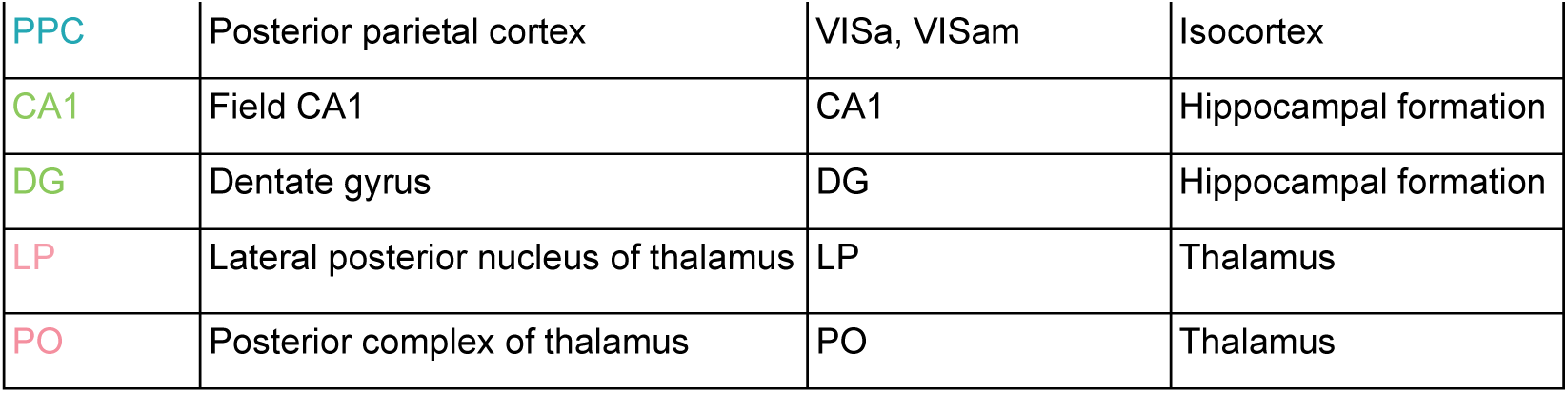
ROI definitions.

The final dataset included 149 mice, 367 recording sessions, and 503 Neuropixels insertions, with up to two insertions per session. Recordings spanned 17 brain regions of interest (Figure 1d; Table 1), including 7 cortical regions, along with structures from the hippocampus, thalamus, midbrain, basal ganglia, and olfactory areas. We applied a series of standardized quality control metrics to the neural data (see Methods).

To quantify pairwise shared variability, we focused on noise correlations (*r_noise_*; Figure 2c). Noise correlations capture the extent to which trial-to-trial fluctuations in firing rate are shared between pairs of neurons (Averbeck et al., 2006; Cohen & Kohn, 2011; Zohary et al., 1994). This measure is widely used to characterize shared variability within neural populations and is directly relevant for understanding how shared fluctuations may shape population coding (Averbeck et al., 2006; Cohen & Kohn, 2011). Here, noise correlations allowed us to test whether ageing alters shared trial-to-trial variability across brain regions and task epochs.

**Figure 2.**
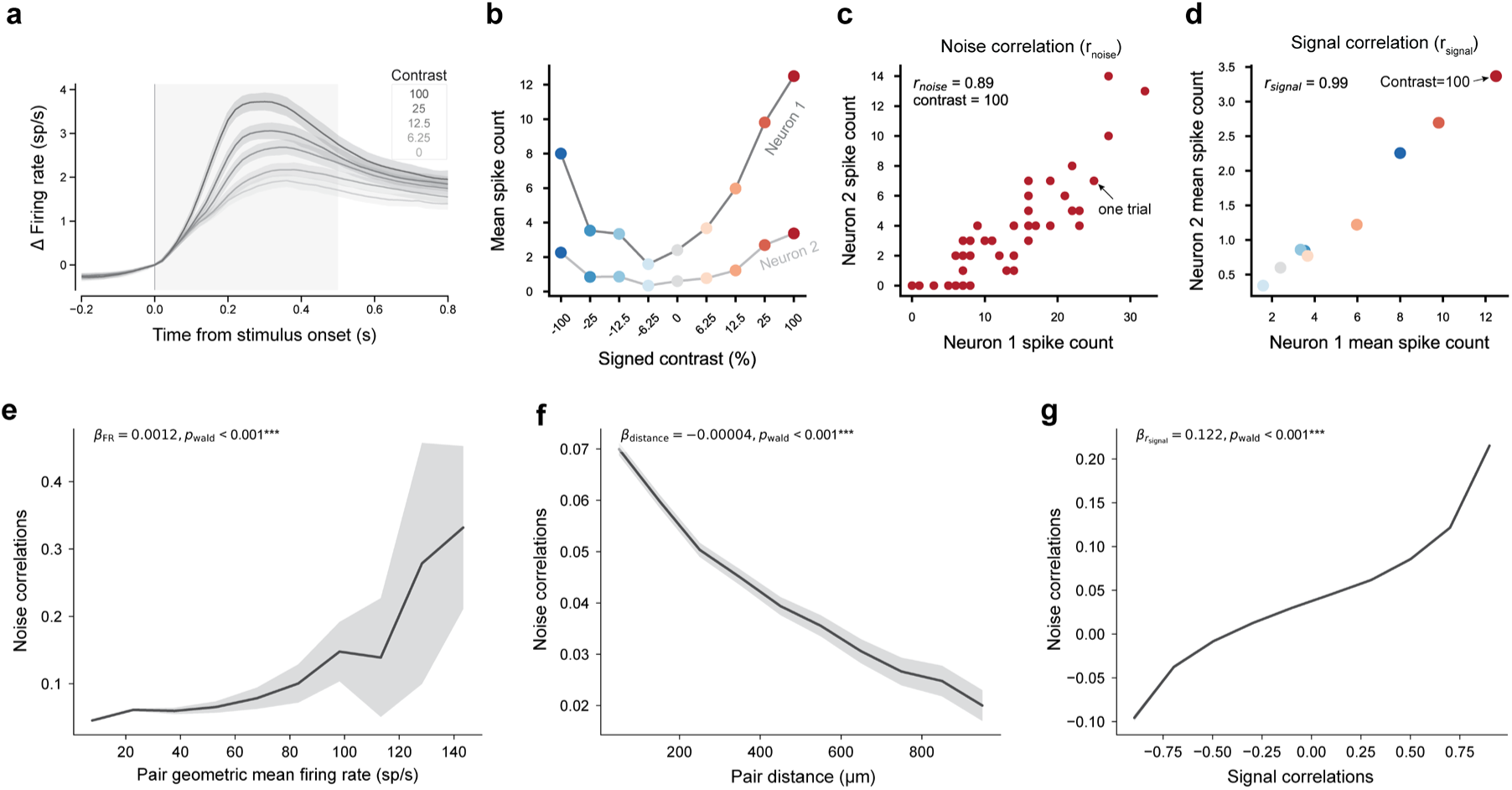
Schematics and validation of metrics. **(a)** Baseline corrected average firing rate time courses across stimulus contrast levels, aligned to stimulus onset. Saturation represents different contrast levels; shaded areas indicate 95% confidence intervals (CIs). The grey area indicates the post-stimulus (0 ms-500 ms) time window used for noise correlations and signal correlations calculation. **(b)** Two example neurons included in panel (a). Mean spike counts within the 0–500 ms post-stimulus window are shown across trials for different signed contrast levels. Different colours represent different signed contrast conditions. **(c)** Schematic definition of noise correlations. Here we use the two example neurons from panel (b) and one contrast level (signed contrast = 100%) for illustration. Each point represents the responses of the two neurons on one trial. Noise correlation, also referred to as spike-count correlation, was measured as the Pearson correlation between trial-to-trial fluctuations in responses to repeated presentations of the same stimulus. **(d)** Schematic definition of signal correlation. Each dot represents the trial-averaged response to one signed contrast level. Signal correlation was measured as the correlation between the two neurons’ mean responses across stimuli. Different colours represent different signed contrast conditions, with the same convention as used in (b). **(e)** The relationship between noise correlations and the geometric mean firing rate of neuron pairs. **(f)** The relationship between noise correlations and the distance between neuron pairs. **(g)** The relationship between noise correlations and signal correlations. For panels (e-g), lines show binned means and shaded areas indicate 95% CIs across all neuron pairs. The examined effects were estimated using liner mixed-effects models with pair geometric mean firing rate/pair distance/signal correlation as a continuous predictor (see Methods). Stars indicate significance based on p values: *p < 0.01, **p < 0.005, and ***p < 0.001.

After preprocessing and quality control, we computed noise correlations for pairs of simultaneously recorded neurons within the same brain region.

### Noise correlations recapitulate canonical physiological relationships

We first confirmed that estimates of trial-to-trial noise correlations reproduced well-established physiological relationships reported in previous work. Neural responses were aligned to stimulus onset, and noise correlations were computed for pairs of simultaneously recorded neurons using spiking activity in a fixed post-stimulus window (0 to 500 ms relative to stimulus onset; Figure 2a–c; see Methods). We then tested whether the noise correlation estimates reproduced three canonical dependencies reported in the literature: their relationships with firing rate, inter-neuronal distance, and signal correlation, a measure of stimulus-tuning similarity between neurons (Figure 2d). We first assessed each relationship in the pooled dataset by comparing a model containing the predictor with a corresponding baseline model without it. These models were fitted to all neuron pairs across brain regions. For all three relationships, both AIC and BIC favoured models containing the predictor over corresponding models without it (see Methods; Supplementary Table S1).

First, consistent with prior work, noise correlations increased with the geometric mean firing rate of the paired neurons (*β*_fr_ = 0.0012, *p*_wald_ < 0.001; ΔAIC = 1423.96; ΔBIC = 1413.20; Figure 2e). This dependence is expected because measured spike-count correlations are sensitive to response strength: when firing rates are low, shared fluctuations are more weakly expressed in observed spike counts, whereas stronger responses make shared variability easier to detect (Cohen & Kohn, 2011).

Second, noise correlations also showed clear spatial organization, with the highest values observed for nearby neuron pairs and progressively lower values at greater inter-neuronal distances (*β*_distance_ = −0.00004, *p*_wald_ < 0.001; ΔAIC = 2191.01; ΔBIC = 2180.24; Figure 2f). This pattern is consistent with the idea that nearby neurons are more likely to share local inputs and circuit interactions. This pattern closely matches classic observations in macaque V1 (Smith & Kohn, 2008; Wang et al., 2019) and in mouse visual cortex (Montijn et al., 2014).

Third, noise correlations were positively related to signal correlation (*β*_r_signal_ = 0.122, *p*_wald_ < 0.001; ΔAIC = 48995.33; ΔBIC = 48984.58; Figure 2g), defined as the similarity between neurons in their stimulus tuning or trial-averaged response profiles (Averbeck et al., 2006; Cohen & Kohn, 2011; Zohary et al., 1994). Thus, neuron pairs with more similar response profiles also tended to share greater trial-to-trial variability, consistent with reports across sensory systems in both mice and primates (Ecker et al., 2014; Montijn et al., 2014; Smith & Kohn, 2008; Smith & Sommer, 2013; Wang et al., 2019). This relationship may arise because similarly tuned neurons are more likely to receive overlapping feedforward, recurrent, or feedback drive (Cohen & Kohn, 2011; Smith & Kohn, 2008).

We next assessed regional variation using region-specific slopes extracted from models containing predictor-by-region interactions. Models including predictor-by-region interactions were favoured over the corresponding common-slope models for all three predictors (all ΔAIC>0; all ΔBIC>0; Supplementary Table S1), indicating regional variation in effect magnitude. Nevertheless, the expected directions were broadly conserved across regions (Supplementary Figures S1–S3). After correcting for multiple comparisons using FDR_0.01_, the expected relationship was significant in 14 of 17 regions for firing rate, 15 of 17 regions for distance, and all 17 regions for signal correlation; LP showed a significant firing-rate relationship in the opposite direction.

Together, these pooled and regional analyses show that our noise-correlation estimates captured established relationships with firing rate, inter-neuronal distance, and signal correlation, supporting their use in the age-related analyses that follow.

### Age effects on noise correlations are region-specific and bidirectional

Following validation of the noise correlation metric, we examined whether ageing was associated with changes in noise correlations. Previous studies have suggested that ageing can increase noise correlations, as reported in primary visual cortex in aged rhesus monkeys (Wang et al., 2019) and mice (Liu et al., 2025), and in auditory cortex in aged mice during passive listening or auditory detection tasks (Shilling-Scrivo et al., 2021, 2022). These findings create an expectation that ageing should increase noise correlations, particularly in sensory cortical populations. At the same time, because these studies focused on specific sensory areas, it remains unclear whether this pattern generalizes across broader cortical and subcortical circuits during active decision-making.

For the main multi-region analyses, we quantified noise correlations between simultaneously recorded neuron pairs within brain region and hemisphere across 17 cortical and subcortical areas (Table 1). Because trials were aligned to stimulus onset, we computed noise correlations separately in matched pre-stimulus (−500 to 0 ms) and post-stimulus (0 to 500 ms) windows, allowing us to examine shared variability before and during stimulus-evoked activity. We first assessed age effects pooled across all recorded brain regions, and then resolved these effects by region.

During the pre-stimulus period, adding a common age effect did not improve model fit relative to the model without age (ΔAIC = −1.66; ΔBIC = −12.42; *β*_age_ = −0.004, *p*_wald_ = 0.558). In contrast, allowing age slopes to vary across regions was favoured over the model without age by both AIC and BIC (ΔAIC = 191.72; ΔBIC = 8.75), indicating substantial regional heterogeneity. Region-resolved analyses revealed that age effects were region-specific and differed in their directionality (Figure 3c,d). Age-related increases were observed in primary visual cortex (VISp; *β*_age_ = 0.096), and the lateral posterior thalamic nucleus (LP; *β*_age_ = 0.029). Age-related decreases were observed in posteromedial visual cortex (VISpm; *β*_age_ = −0.078), motor related cortex (MOs; *β*_age_ = −0.046) and motor-related superior colliculus (SCm; *β*_age_ = −0.051; all *p*_fdr_ < 0.01).

**Figure 3.**
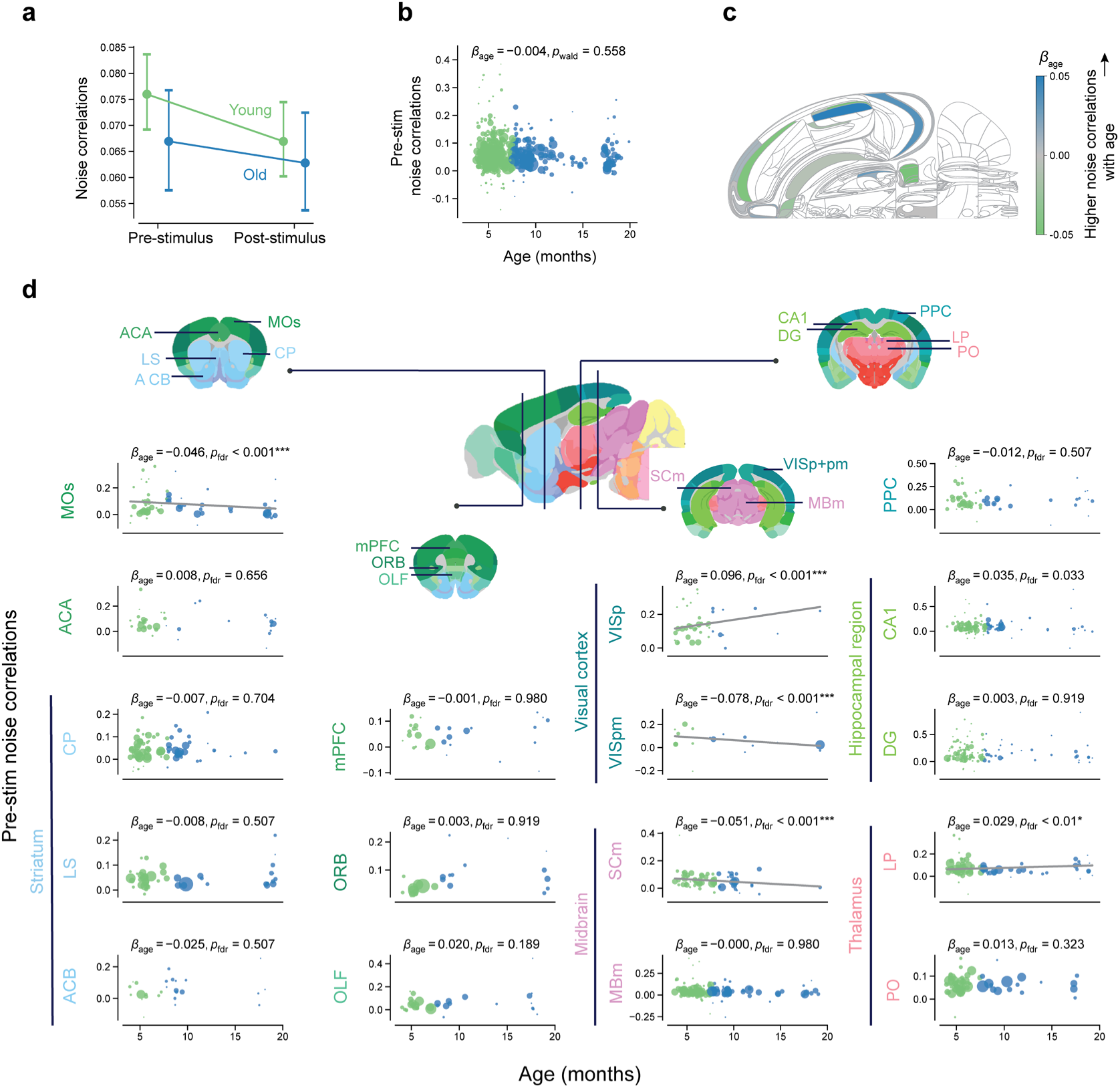
Age effects on pre-stimulus noise correlations. **(a)** Pooled noise correlations across neuron pairs, summarized separately for each age group and time window. Neuron-pair correlations were first averaged within each recording session, then across sessions. Error bars show 95% CIs. **(b)** The relationship between pre-stimulus noise correlations and mouse age. Each dot represents one insertion, with dot size indicating the number of neuron pairs included in the insertion-level estimate. **(c)** The Swanson map showing region-specific age effects on pre-stimulus noise correlations. The colour indicates the estimated slope of age. White areas were not covered in our recordings. **(d)** Regional specificity of the relationship between pre-stim noise correlations and mouse age. Each dot represents the mean noise correlation for one insertion. The grey line visualizes the age slope estimated from the age-by-region interaction model (see Methods). Lines are drawn only for regions whose age slope has an FDR-adjusted p value < 0.001. Annotations report the estimated age coefficient and its FDR-adjusted p value. Stars indicate significance based on FDR-adjusted p values: *p < 0.01, **p < 0.005, and ***p < 0.001.

Thus, ageing did not produce a change in pre-stimulus noise correlations that was shared across brain areas. Instead, noise correlations show spatially heterogeneous age effects, with some areas increasing and others decreasing in noise correlations in older animals.

Post-stimulus noise correlations in the matched 0–500-ms window showed a broadly similar pattern (Supplementary Figure S4c,d). There was again no evidence for an age effect shared across regions (ΔAIC = −1.98; ΔBIC = −12.74; *β*_age_ = −0.001, *p*_wald_ = 0.893; Supplementary Figure S4b). AIC favoured allowing age slopes to vary by region, whereas BIC modestly favoured the model without age (ΔAIC = 179.13; ΔBIC = −3.84). Nevertheless, the region-specific estimates reproduced several features of the pre-stimulus pattern: age was positively associated with noise correlations in VISp (*β*_age_ = 0.140) but negatively associated with noise correlations in MOs (*β*_age_ = −0.040) and VISpm (*β*_age_ = −0.084; all *p*_fdr_ < 0.01; Supplementary Figure S4d). Thus, the post-stimulus results also indicated bidirectional regional age effects, although support for the overall region-specific model differed between AIC and BIC.

The positive age effect in primary visual cortex was of particular relevance to previous work. Specifically, in a recent study, Liu et al. (2025) measured noise correlations between primary visual cortical neurons while mice passively viewed Gabor stimuli, comparing young (2 months) and aged (12 months) animals. The recording targets from Liu et al. were located within the VISp region analysed here (Figure 4a), providing an anatomical basis for comparison. To further align our analysis with theirs, we repeated the analysis using their post-stimulus window of 0– 1000 ms. Noise correlations in VISp increased with age (*β*_age_ = 0.176, *p*_fdr_ < 0.001; Figure 4b). This is consistent with the effect reported by Liu et al. despite differences in task context, behavioural state, and dataset. The complete 1000-ms regional pattern is presented in Supplementary Figure S5 and showed that the positive VISp effect formed part of a broader, bidirectional pattern across brain regions.

**Figure 4.**
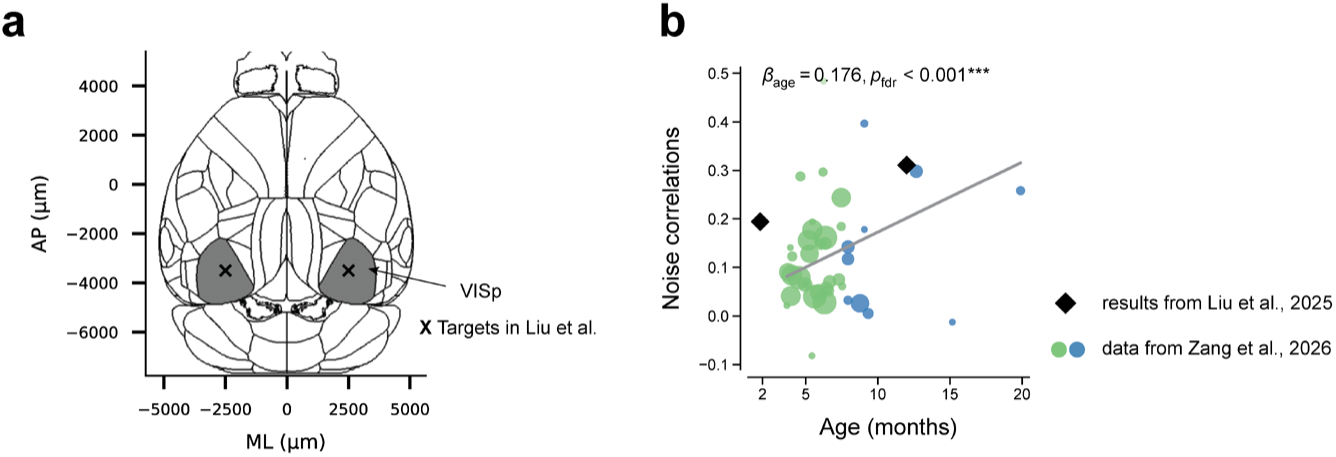
Replication of previous findings in mouse primary visual cortex. **(a)** Crosses indicate the target locations used by Liu et al. (2025) to record neural activity from mouse primary visual cortex (V1). Grey regions indicate VISp, the primary visual cortex region defined in the IBL Beryl atlas. The Liu et al. recording targets fall within the VISp region analysed here, supporting anatomical comparison between the two datasets. **(b)** Relationship between stimulus-period noise correlation and mouse age in primary visual cortex. To enable comparison with Liu et al. (2025), this analysis used a 0–1000 ms window after stimulus onset. Black diamonds show the group-level median noise correlation values reported by Liu et al. (2025) for their young and aged groups. Coloured dots represent median noise correlations values from one insertion, with dot size indicating the number of neuron pairs included in the insertion-level estimate. The grey line visualizes the age slope estimated from the mixed-effects model. Stars indicate significance based on FDR-adjusted p values: *p < 0.01, **p < 0.005, and ***p < 0.001.

Together, these findings help reconcile the absence of a brain-wide age effect with previous single-region studies reporting elevated noise correlations with ageing in the sensory cortex (Shilling-Scrivo et al., 2021, 2022; Liu et al., 2025; Wang et al., 2019). The VISp-specific increase in noise correlations did not generalize across regions. Instead, it coexisted with bidirectional age effects across cortical and subcortical regions, indicating that single-region observations from sensory cortex may capture local age effects without fully reflecting the broader regional organization of noise correlations.

### Stimulus-induced reductions in noise correlations show limited age-related differences

Stimulus onset is known to reduce single-neuron trial-to-trial spike-count variability, commonly quantified by the Fano factor, a phenomenon termed variability quenching that has been observed across multiple cortical regions and species (Churchland et al., 2010; Licata et al., 2017; Poland et al., 2019). Stimulus-related reductions in noise correlations have also been reported in primate visual cortex and rodent olfactory cortex (Iurilli & Datta, 2017; Miura et al., 2012; Smith & Kohn, 2008). To first establish whether noise correlations were modulated by stimulus onset in the present dataset, we quantified Δnoise correlations as post-stimulus minus pre-stimulus noise correlations. Across regions, noise correlations showed a modest overall reduction after stimulus onset (adjusted mean Δnoise correlations = −0.007, *p*_wald_ < 0.001; Supplementary Figure S6a). Given our previous finding that Fano factor quenching was attenuated in older mice (Zang et al., 2026), we then asked whether the magnitude of this stimulus-related reduction in noise correlations differed with age.

This reduction showed no age-related change shared across regions: adding a common age effect did not improve model fit relative to the model without age (ΔAIC = −1.10; ΔBIC = −11.86; *β*_age_ = 0.004, *p*_wald_ = 0.342; Supplementary Figure S6b), providing no evidence for an age-related change shared across regions. Model-selection criteria provided conflicting evidence for regional differences: AIC favoured allowing age slopes to vary across regions, whereas BIC favoured the model without any age term (ΔAIC = 66.43; ΔBIC = −116.54). Within the region-specific model, positive age slopes survived multi-comparison correction (FDR_0.01_) in the nucleus accumbens (ACB; *β*_age_ = 0.095, *p*_fdr_ < 0.001) and posterior parietal cortex (PPC; *β*_age_ = 0.042, *p*_fdr_ < 0.001; Supplementary Figure S6c,d), consistent with weaker stimulus-related reductions in these regions. Thus, evidence for age-related differences in stimulus-induced noise-correlation modulation was restricted to specific regions and depended on the model-selection criterion.

## Discussion

In this study, we characterized age-related differences in shared neural variability during visual decision-making. Across 348,937 simultaneously recorded within-region neuron pairs from 17 brain regions, the principal finding was regional heterogeneity: age associations with pre- and post-stimulus noise correlations differed in direction across regions. Within this pattern, the age-related increase in primary visual cortex replicated previous findings, showing that a reproducible sensory-cortical effect can coexist with opposing age associations elsewhere. Evidence for age-related differences in stimulus-induced changes was more limited and depended on the model-selection criterion. Together, these findings indicate that age-related differences in shared variability form a regionally heterogeneous pattern rather than reflecting a brain-wide shift. They should therefore be interpreted at a region-resolved level rather than inferred from a single brain area.

These region-resolved results place previous reports of increased noise correlations in ageing sensory cortex in a broader context (Liu et al., 2025; Shilling-Scrivo et al., 2021, 2022; Wang et al., 2019).The replication in primary visual cortex indicates that this previously reported effect can be reproduced in an independent dataset and during active decision-making. However, the divergence between primary and higher visual cortex suggests that this effect is not a general property of all visually responsive regions in the mouse brain. By comparing cortical and subcortical regions within the same behavioural paradigm and analytical framework, the present study identifies regional specificity as a central feature of age-related differences in shared variability, highlighting the need for caution when extrapolating from single-region studies to broader theories of age-related neural change (Ryan et al., 2026).

The regional specificity of these age effects raises the possibility that ageing-related changes in shared variability are shaped by the circuit architecture of each brain region. Ageing can alter cortical excitatory and inhibitory connectivity (Xue et al., 2023), while perturbing different inhibitory interneuron subtypes can have opposing effects on noise correlations (Xu et al., 2026). Cholinergic modulation can also influence these correlations (Minces et al., 2017). Together with regional differences in interneuron composition (Kim et al., 2017), these findings suggest that age-related changes in inhibition or neuromodulatory drive may have different consequences for shared variability across circuits.

Evidence for age-related differences in stimulus-induced modulation was more limited. Stimulus onset is widely documented to quench single-neuron trial-to-trial variability, marking a transition from ongoing activity to a more stimulus-constrained processing state (Churchland et al., 2010; Ponce-Alvarez et al., 2013). Noise correlations also decreased modestly after stimulus onset in the present dataset. However, within the region-specific model, positive age associations with this change were supported only in ACB and PPC, and evidence for age-by-region variation differed between model-selection criteria. Moreover, the regional profile did not fully overlap with that of age-related differences in Fano factor quenching observed previously in the same datasets (Zang et al., 2026). Fano factors quantify the trial-to-trial variability of individual neurons, whereas noise correlations quantify normalized co-fluctuations between neurons. This contrast also suggests that age-related changes in single-neuron variability do not necessarily translate into parallel changes in variability shared across neuronal populations.

It is not trivial to determine the functional role of noise correlation reductions in neural coding. The impact of shared variability on population coding and behaviour depends on its structure, particularly whether correlated fluctuations align with stimulus-, choice-, or movement-related coding axes, rather than on the average pairwise correlation alone (Averbeck et al., 2006; Moreno-Bote et al., 2014; Kohn et al., 2016). For this reason, the regional increases or decreases observed here do not by themselves imply better or worse neural coding or behavioural performance. This interpretive caution is particularly important in the present study because our recordings were obtained during active visual decision-making, where attention, arousal, locomotion, and task engagement can modulate correlated variability (Cohen & Maunsell, 2009; McGinley et al., 2015; Shi et al., 2022; Vinck et al., 2015). Moreover, because wheel movements controlled stimulus position, trial-to-trial differences in movement during the post-stimulus period may have contributed to correlated neural fluctuations that were not accounted for by stimulus contrast alone. Accordingly, the post-stimulus associations may reflect task-related shared variability rather than purely sensory-driven correlations. These considerations constrain the functional attribution of the observed associations but do not alter the central finding that their anatomical distribution is regionally heterogeneous.

In summary, this study provides a large-scale, region-resolved characterization of age-related differences in pairwise shared neural variability during visual decision-making. Across the sampled cortical and subcortical regions, age associations were heterogeneous and bidirectional rather than consistent with a brain-wide shift, whereas associations with stimulus-induced change were more spatially restricted. By placing the reproducible VISp increase within this broader pattern, our findings show that previous sensory-cortex observations do not generalize uniformly across the sampled brain regions. The partially overlapping regional profiles of age associations with noise-correlation modulation and Fano factor quenching further suggest that single-neuron and shared variability need not show parallel age-related differences. Together, these findings identify regional specificity as a central feature of age-related differences in shared neural variability during visual decision-making.

## Methods

### Data, animals, and the behavioural task

This study builds on our previous analysis of age-related changes in single-neuron variability in the IBL visual decision-making task (Zang et al., 2026). The same dataset and preprocessing pipeline were used unless otherwise specified. We included 130 mice from a previously released dataset (C57BL/6, 89 male, mean age = 6.64 months, range 3.10–15.13 months) and 19 mice from a newly recorded dataset (C57BL/6, 11 male, mean age = 16.58 months, range 10.58–19.90 months). After quality control, 149 mice were included in the formal analysis. For visualization, mice were categorized based on their age at recording into a young group (N = 97, mean age 5.50 months) and an old group (N = 52, mean age 11.56 months), using 7.6 months (mean age in the dataset) as the age cutoff.

Mice were trained to perform a standardized visual decision-making task (International Brain Laboratory et al., 2021). In this task, mice had to decide the location of a visual stimulus presented on the screen in front of them. Specifically, each trial began when the mouse held the wheel still for 0.4-0.7 seconds. Then, an auditory cue (a 100-ms tone, 5 kHz sine wave) and a visual stimulus (Gabor patch) were presented on either the left or right side of the screen. Mice indicated the stimulus location by turning the response wheel to bring the stimulus to the centre of the screen. The contrast level of the visual stimulus varied across trials. There were five different contrast levels (%): 100, 25, 12.5, 6.25, 0. Combining the five contrast levels with the two stimulus sides (left, right) resulted in nine conditions: −100, −25, −12.5, −6.25, 0, 6.25, 12.5, 25, 100 (positive for right stimuli, negative for left stimuli). Correct responses were rewarded with sugar water, while incorrect responses were followed by a noise burst and a longer inter-trial interval. The behavioural task is further described in (International Brain Laboratory et al., 2021). The detailed protocol for animal training can be found in the Methods section of (International Brain Laboratory et al., 2021) and the IBL protocol for mice training (International Brain Laboratory, 2020).

Extracellular recordings were acquired using Neuropixels probes (Jun et al., 2017). For a detailed description of the animal surgery, apparatus, and recording procedure, see Appendix 2 and 3 in (International Brain Laboratory, Banga, et al., 2025). Up to two probes were inserted per recording session.

Data preprocessing followed the procedures described previously (Zang et al., 2026), including sessions and trials filtering, spike sorting, and neuron quality control.

We included 17 brain regions (in cortex, striatum, midbrain, hippocampus, and thalamus; Table 1). These regions were selected based on a combination of scientific, statistical, and practical considerations: scientifically, frontal cortical areas are known to show early age-related decline; statistically, the set includes the ‘repeated site’ used to assess cross-laboratory reproducibility (International Brain Laboratory, Banga, et al., 2025); and practically, these areas offered reliable surgical accessibility.

### Noise correlations and signal correlations

We computed noise correlations for pairs of neurons recorded within the same brain region and from the same Neuropixels probe to ensure anatomical consistency and recording simultaneity. Noise correlations were computed following standard practice in previous literature (Cohen & Kohn, 2011; Ecker et al., 2014; Ruff & Cohen, 2014). Specifically, for each stimulus condition, we counted spikes of each neuron in a fixed analysis window (500-ms window either before or after stimulus onset, depending on the analysis) in each trial. We then took the Z-scored responses and concatenated all trials for each neuron, and the Pearson correlation coefficient was then computed for each pair of neurons within the same brain region, recorded from the same probe.

Signal correlations were computed following standard definitions (Cohen & Kohn, 2011; Ecker et al., 2010; Ruff & Cohen, 2016). For each stimulus condition, we calculated the mean spike count of each neuron in a fixed analysis window (0-500 ms) relative to stimulus onset. The vector of mean responses across conditions defined the neuron’s tuning curve. For each simultaneously recorded pair of neurons, the signal correlation was then defined as the Pearson correlation coefficient between their tuning curves (i.e., between condition-averaged neural responses across all 9 included conditions).

### Neuron pair distance

The distance between a pair of neurons is defined as the difference in depth of the two channels on the Neuropixel probe where the pair was recorded.

### Firing rate

Because noise correlation estimates can depend on firing rate, we calculated the geometric mean firing rate of each neuron pair and included it as a covariate in all models of age effects. This regression-based adjustment allowed age to remain a continuous predictor while retaining the full sample.

### Statistical tests

Statistical analyses of pairwise noise correlations were conducted using linear mixed-effects models fitted at the neuron-pair level. To account for the non-independence of neuron pairs recorded from the same session, all models included a random intercept for recording sessions. For all analyses, we specified a set of nested candidate models representing increasingly complex hypotheses. Relative model support was assessed using both the Akaike information criterion (AIC) and the Bayesian information criterion (BIC), which applies a stronger penalty for model complexity. Results from both criteria were reported, and disagreements in model support were explicitly noted. We defined ΔAIC and ΔBIC as the value for the simpler model minus that for the more complex model; therefore, positive values favour the more complex model and negative values favour the simpler model.

#### Validation of noise correlations

We first validated the expected relationships between noise correlations and pair mean firing rate, pair distance, and signal correlation. For each predictor, we fitted the following models:

`M0: r_noise ∼ region + (1 | session)`
`M_predictor: r_noise ∼ predictor + region + (1 | session)`
`M_interaction: r_noise ∼ predictor × region + (1 | session)`

The comparison between M_predictor and M0 tested for a common association between the predictor and noise correlations across regions. The comparison between M_interaction and M_predictor tested whether its associations differed among regions, whereas the comparison between M_interaction and M0 tested whether the data supported any predictor-related structure.

For the pooled validation analyses, we report the common predictor coefficient and its two-sided Wald p-value from M_predictor. Region-specific predictor slopes were estimated from the corresponding joint interaction model. Their Wald p-values were adjusted across the 17 regions separately for each validation relationship using the Benjamini–Hochberg false-discovery-rate (BH-FDR) procedure, with (q<.01) used as the significance threshold.

#### Age effects on noise correlations

For each analysis metric—pre-stimulus noise correlations, post-stimulus noise correlations, and Δnoise correlations—we fitted three prespecified models:

`M0: metric ∼ region + covariates + (1 | session)`
`M_age: metric∼ age + region + covariates + (1 | session)`
`M_interaction: metric ∼ age × region + covariates + (1 | session)`

Here, the covariates were the number of trials, pair geometric mean firing rate, and neuron pair distance. The comparison between M_age and M0 tested whether there was a common age association across regions. The comparison between M_interaction and M_age tested whether age associations differed among regions, whereas the comparison between M_interaction and M0 tested whether the data supported any age-related structure.

When the interaction model was supported, region-specific age slopes were estimated from the joint M_interaction model. For each region, we report the age coefficient *β*_age_, and its two-sided Wald p-value. The regional p-values were adjusted across the 17 regions separately for each analysis metric using the BH-FDR procedure, with (q<.01) used as the significance threshold.

## Data and code availability

The data analysed in this study are publicly available. Instructions for accessing these data are provided in the IBL Library (https://docs.internationalbrainlab.org/notebooks_external/2025_data_release_aging_zang.html), and GitHub repo (https://github.com/Fenying-Zang/Ageing_behavioral_and_neural_variability).

Data analyses were performed using custom Python scripts in a reproducible Python environment. All code required to reproduce the figures and analyses is available at https://github.com/Fenying-Zang/ageing-noise-correlations.

## Acknowledgements

We thank the CoCoSys lab for helpful discussions and feedback, and the International Brain Laboratory for support with data analysis. We are grateful to Sander Nieuwenhuis for his helpful comments on the first draft and Steven Mileti c for advice on statistical modelling.

## Funding

AEU was supported by the German National Academy of Sciences Leopoldina, the International Brain Research Organization, and a Veni fellowship (VI.Veni.212.184) from the Netherlands Organisation for Scientific Research. FZ was supported by a PhD fellowship (202204910080) from the Chinese Scholarship Council.

## Author contributions

Conceptualization: F.Z. and A.E.U.; methodology: F.Z. and A.E.U.; formal analysis: F.Z.; data curation: F.Z.; visualization: F.Z. and A.E.U.; writing: F.Z. and A.E.U.; funding acquisition: F.Z. and A.E.U.; project administration: F.Z. and A.E.U.; supervision: A.E.U.

## Competing interests

The authors declare no competing interests.

## Supplementary figures

**Figure S1.**
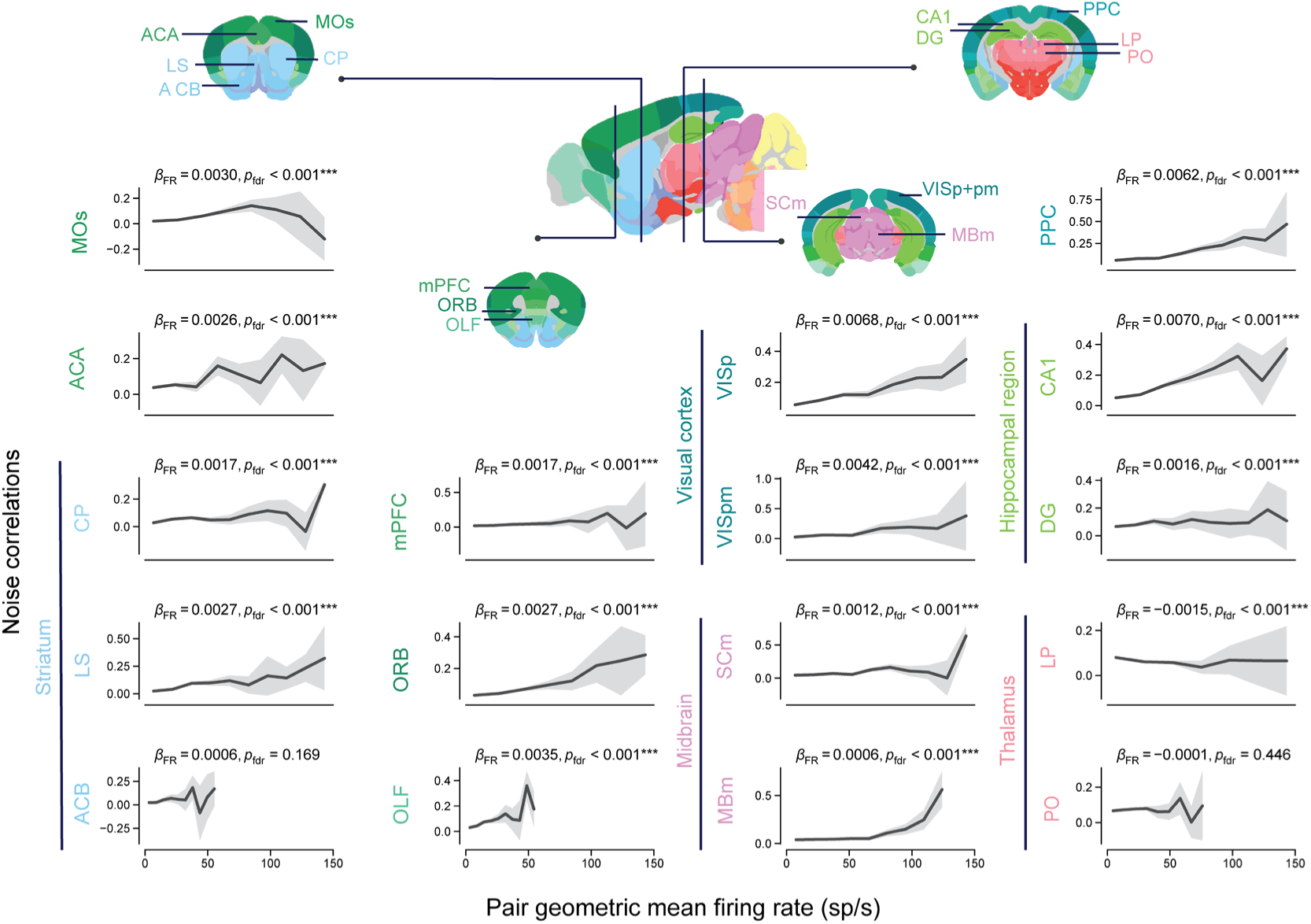
The relationship between noise correlations and the geometric mean firing rate of neuron pairs in each region. Lines show binned means calculated within each region, and shaded areas indicate 95% CIs for these means. Region-specific firing-rate slopes were estimated from a linear mixed-effects model including an interaction between pair geometric mean firing rate and brain region (see Methods). Annotations report the estimated regional slope and its FDR-adjusted p value. Stars indicate significance of the regional slopes based on FDR-adjusted p values: *p < 0.01, **p < 0.005, and ***p < 0.001.

**Figure S2.**
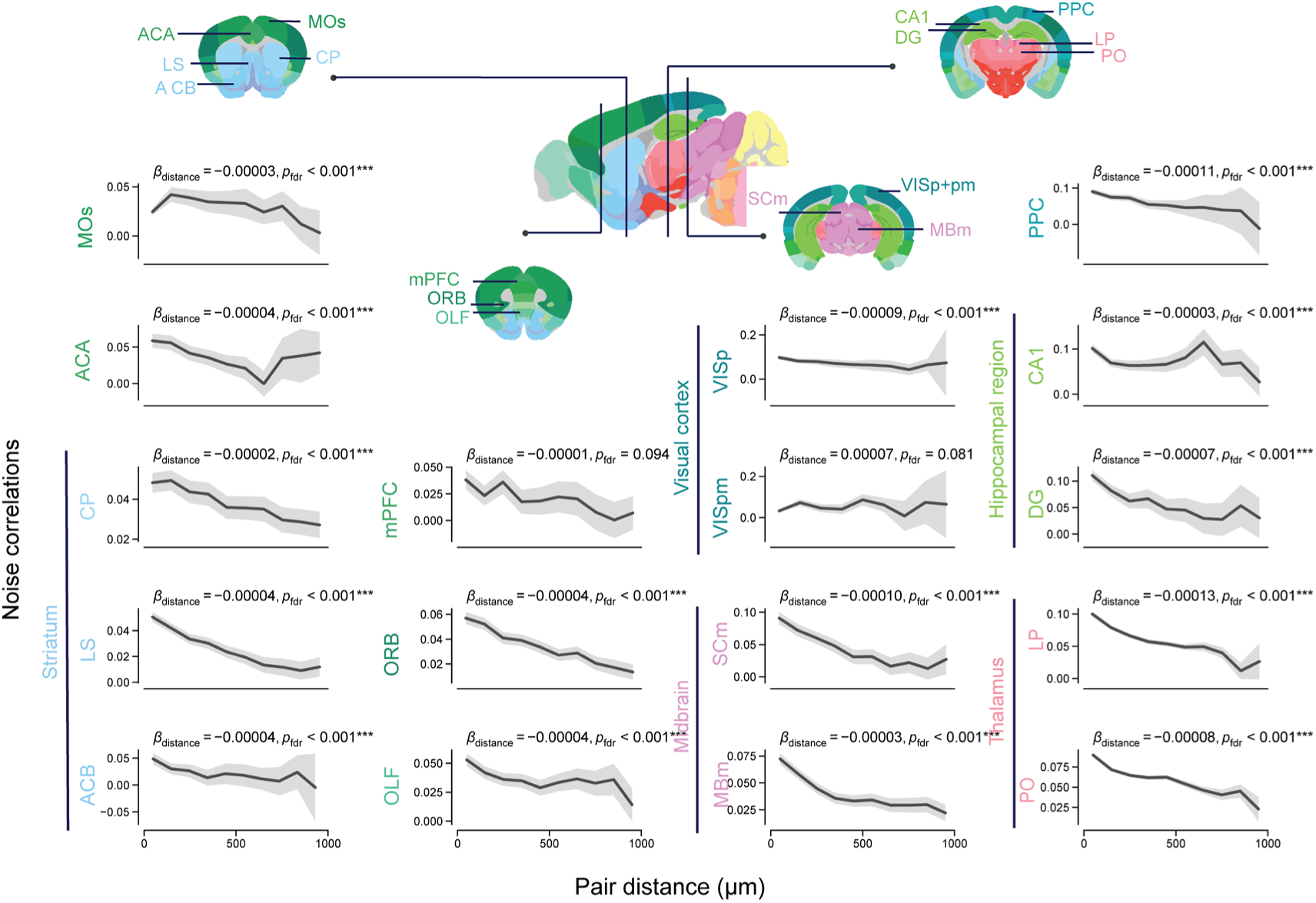
The relationship between noise correlations and the distance between the two neurons within each pair, shown separately for each region. Lines show binned means calculated within each region, and shaded areas indicate 95% CIs for these means. Region-specific distance slopes were estimated from a linear mixed-effects model including an interaction between pair distance and brain region (see Methods). Annotations report the estimated regional slope and its FDR-adjusted p value. Stars indicate significance of the regional slopes based on FDR-adjusted p values: *p < 0.01, **p < 0.005, and ***p < 0.001.

**Figure S3.**
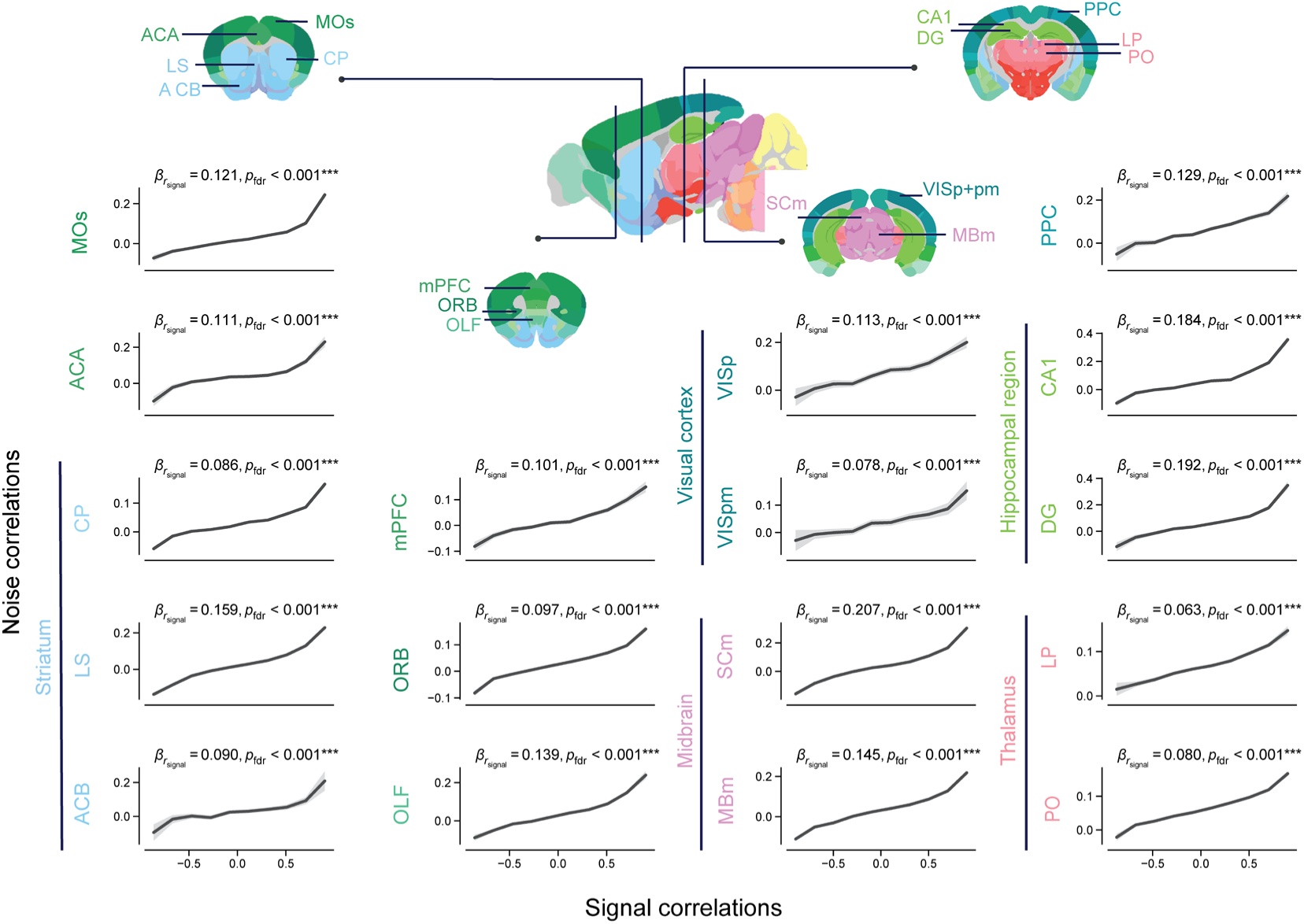
The relationship between noise correlations and signal correlations in each region. Lines show binned means calculated within each region, and shaded areas indicate 95% CIs for these means. Region-specific signal-correlation slopes were estimated from a linear mixed-effects model including an interaction between signal correlation and brain region (see Methods). Annotations report the estimated regional slope and its FDR-adjusted p value. Stars indicate significance of the regional slopes based on FDR-adjusted p values: *p < 0.01, **p < 0.005, and ***p < 0.001.

**Figure S4.**
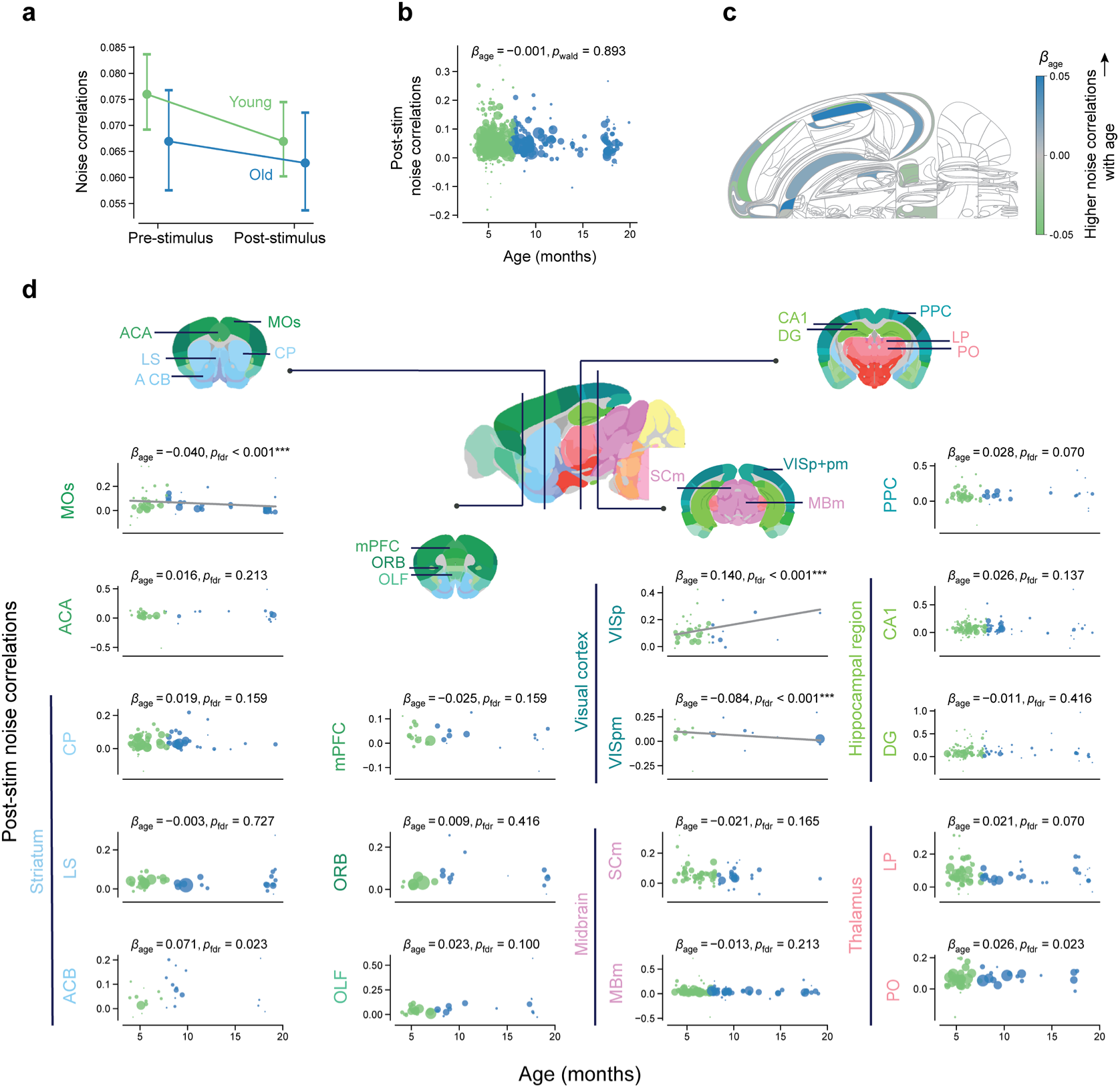
Age effects on post-stimulus (500 ms window) noise correlations. **(a)** Pooled noise correlations across neuron pairs, summarized separately for each age group and time window. Neuron-pair correlations were first averaged within each recording session, then across sessions. Error bars show 95% CIs. **(b)** The relationship between post-stimulus noise correlations and mouse age. Each dot represents one insertion, with dot size indicating the number of neuron pairs included in the insertion-level estimate. **(c)** The Swanson map showing region-specific age effects on post-stimulus noise correlations. The colour indicates the estimated slope of age. White areas were not covered in our recordings. **(d)** Regional specificity of the relationship between post-stim noise correlations and mouse age. Each dot represents the mean noise correlation for one insertion. The grey line visualizes the age slope estimated from the age-by-region interaction model (see Methods). Lines are drawn only for regions whose age slope has an FDR-adjusted p value < 0.001. Annotations report the estimated age coefficient and its FDR-adjusted p value. Stars indicate significance based on FDR-adjusted p values: *p < 0.01, **p < 0.005, and ***p < 0.001.

**Figure S5.**
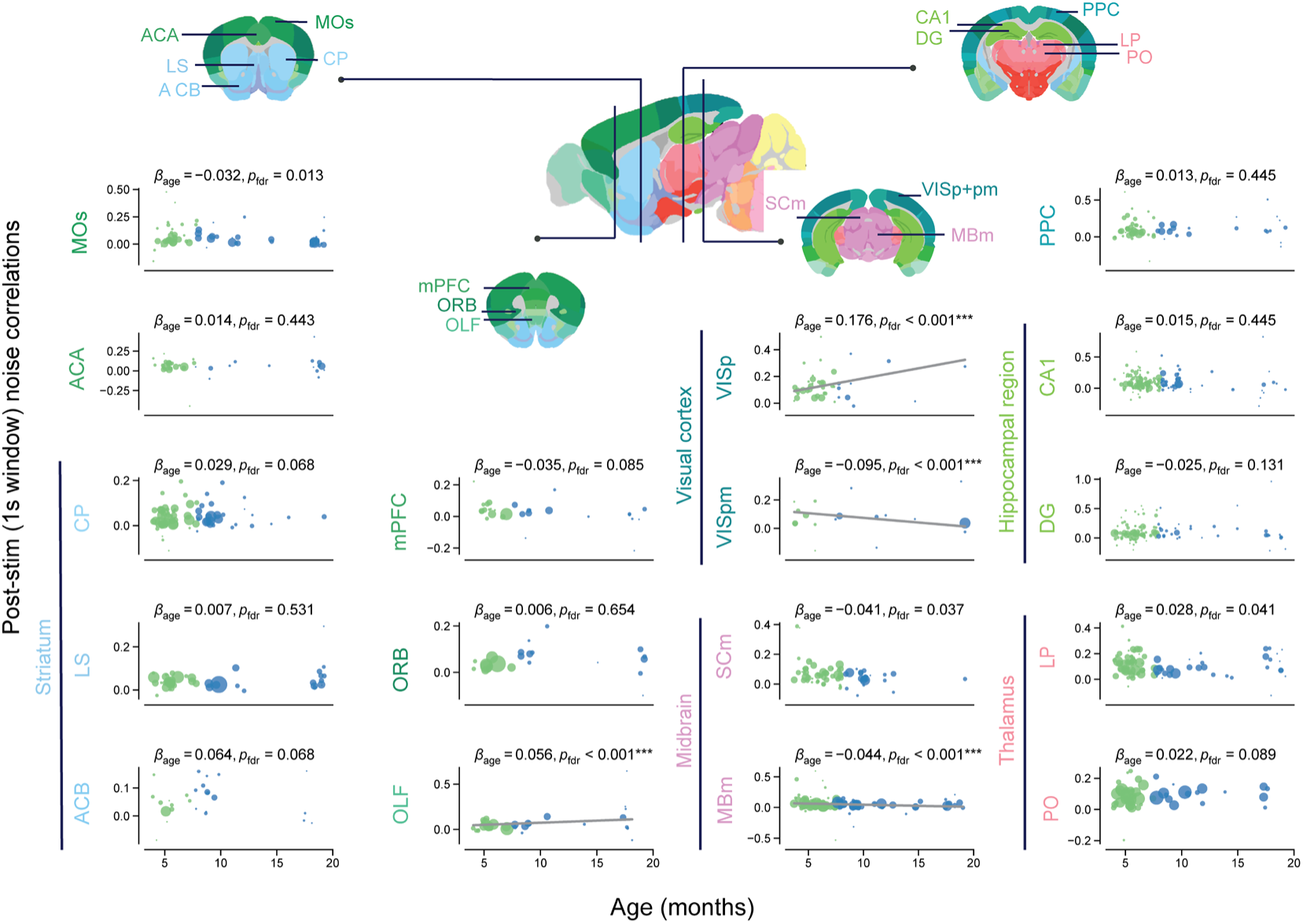
Age effects on post-stimulus (1000ms window) noise correlations. Regional specificity of the relationship between post-stim noise correlations and mouse age. Note: The longer-window analysis again provided no support for an age effect shared across regions (ΔAIC = −1.73; ΔBIC = −12.50), whereas both AIC and BIC favoured allowing age slopes to vary by region (ΔAIC = 190.59; ΔBIC = 7.62). Each dot represents the mean noise correlation for one insertion. The grey line visualizes the age slope estimated from the age-by-region interaction model (see Methods). Lines are drawn only for regions whose age slope has an FDR-adjusted p value < 0.001. Annotations report the estimated age coefficient and its FDR-adjusted p value. Stars indicate significance based on FDR-adjusted p values: *p < 0.01, **p < 0.005, and ***p < 0.001.

**Figure S6.**
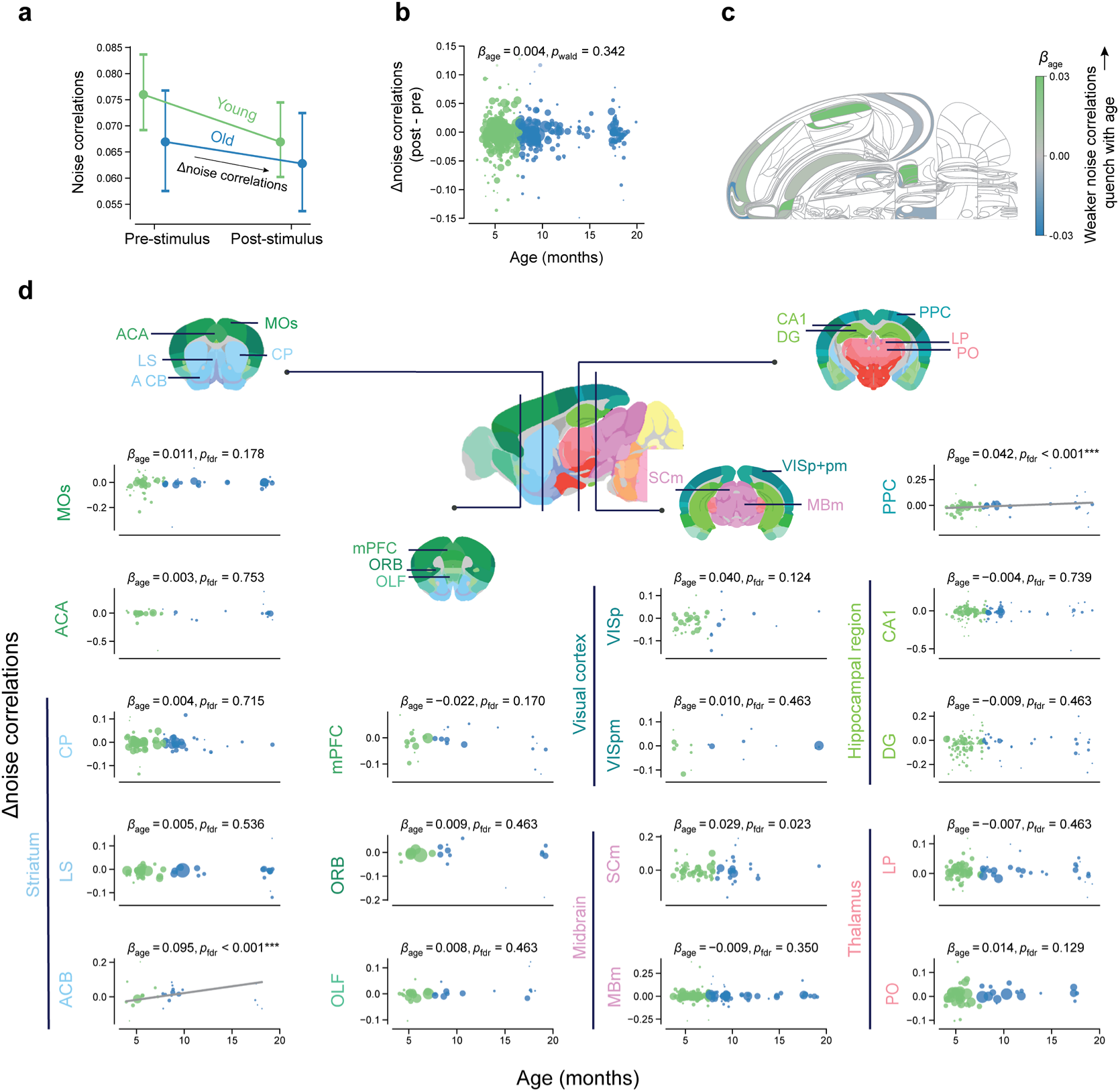
Age effects on Δnoise correlations. **(a)** Pooled noise correlations across neuron pairs, summarized separately for each age group and time window. Neuron-pair correlations were first averaged within each recording session, then across sessions. Error bars show 95% CIs. **(b)** The relationship between Δnoise correlations (computed as post-stimulus noise correlations minus pre-stimulus noise correlations) and mouse age. Each dot represents one insertion, with dot size indicating the number of neuron pairs included in the insertion-level estimate. **(c)** The Swanson map showing region-specific age effects on Δnoise correlations. The colour indicates the estimated slope of age. White areas were not covered in our recordings. **(d)** Regional specificity of the relationship between Δnoise correlations and mouse age. Each dot represents the mean Δnoise correlations for one insertion. The grey line visualizes the age slope estimated from the age-by-region interaction model (see Methods). Lines are drawn only for regions whose age slope has an FDR-adjusted p value < 0.001. Annotations report the estimated age coefficient and its FDR-adjusted p value. Stars indicate significance based on FDR-adjusted p values: *p < 0.01, **p < 0.005, and ***p < 0.001.

## Supplementary Table

**Table S1.**
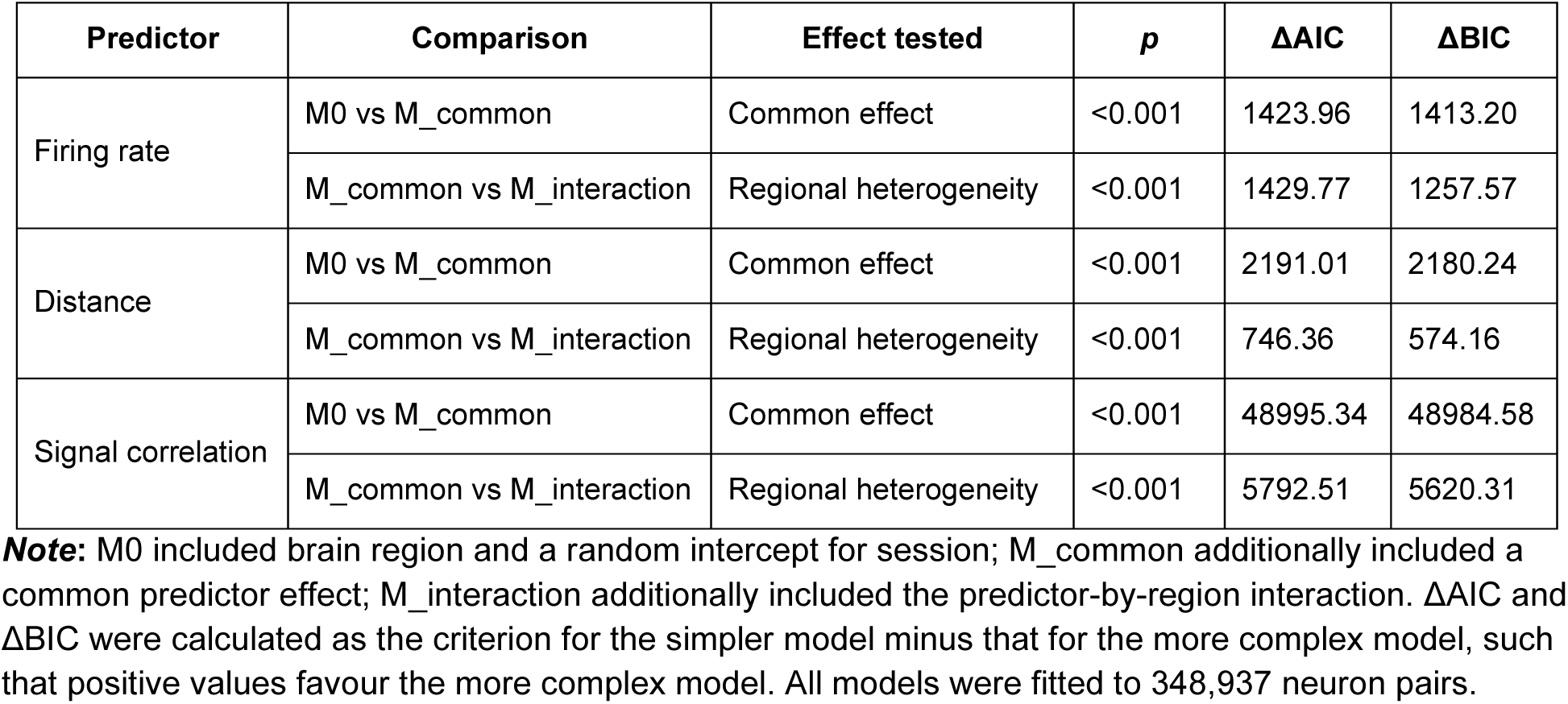
Model comparison results.

## Notes

### Competing Interest Statement

The authors have declared no competing interest.

### Summary of Updates

This version includes additional validation analyses and revised statistical analyses. The figures and text have been updated accordingly.

## References

Averbeck, B. B., Latham, P. E., & Pouget, A. (2006). Neural correlations, population coding and computation. Nature Reviews Neuroscience, 7(5), 358–366. 10.1038/nrn1888

Botvinik-Nezer, R., Holzmeister, F., Camerer, C. F., Dreber, A., Huber, J., Johannesson, M., Kirchler, M., Iwanir, R., Mumford, J. A., Adcock, R. A., Avesani, P., Baczkowski, B. M., Bajracharya, A., Bakst, L., Ball, S., Barilari, M., Bault, N., Beaton, D., Beitner, J., … Schonberg, T. (2020). Variability in the analysis of a single neuroimaging dataset by many teams. Nature, 582(7810), 84–88. 10.1038/s41586-020-2314-9

Churchland, M. M., Yu, B. M., Cunningham, J. P., Sugrue, L. P., Cohen, M. R., Corrado, G. S., Newsome, W. T., Clark, A. M., Hosseini, P., Scott, B. B., Bradley, D. C., Smith, M. A., Kohn, A., Movshon, J. A., Armstrong, K. M., Moore, T., Chang, S. W., Snyder, L. H., Lisberger, S. G., … Shenoy, K. V. (2010). Stimulus onset quenches neural variability: A widespread cortical phenomenon. Nature Neuroscience, 13(3), Article 3. 10.1038/nn.2501

Cohen, M. R., & Kohn, A. (2011). Measuring and interpreting neuronal correlations. Nature Neuroscience, 14(7), Article 7. 10.1038/nn.2842

Cohen, M. R., & Maunsell, J. H. R. (2009). Attention improves performance primarily by reducing interneuronal correlations. Nature Neuroscience, 12(12), Article 12. 10.1038/nn.2439

Cottam, N. C., Ofori, K., Stoll, K. T., Bryant, M., Rogge, J. R., Hekmatyar, K., Sun, J., & Charvet, C. J. (2025). From Circuits to Lifespan: Translating Mouse and Human Timelines with Neuroimaging-Based Tractography. Journal of Neuroscience, 45(12). 10.1523/JNEUROSCI.1429-24.2025

Cremer, R., & Zeef, E. J. (1987). What Kind of Noise Increases With Age? Journal of Gerontology, 42(5), 515–518. 10.1093/geronj/42.5.515

Crossman, E. R. & Szafran, J. (1956). Changes with age in the speed of information-intake and discrimination. Experientia, 128–135.

Doiron, B., Litwin-Kumar, A., Rosenbaum, R., Ocker, G. K., & Josić, K. (2016). The mechanics of state-dependent neural correlations. Nature Neuroscience, 19(3), Article 3. 10.1038/nn.4242

Ecker, A. S., Berens, P., Cotton, R. J., Subramaniyan, M., Denfield, G. H., Cadwell, C. R., Smirnakis, S. M., Bethge, M., & Tolias, A. S. (2014). State Dependence of Noise Correlations in Macaque Primary Visual Cortex. Neuron, 82(1), 235–248. 10.1016/j.neuron.2014.02.006

Ecker, A. S., Berens, P., Keliris, G. A., Bethge, M., Logothetis, N. K., & Tolias, A. S. (2010). Decorrelated Neuronal Firing in Cortical Microcircuits. Science, 327(5965), 584–587. 10.1126/science.1179867

Gu, Y., Liu, S., Fetsch, C. R., Yang, Y., Fok, S., Sunkara, A., DeAngelis, G. C., & Angelaki, D. E. (2011). Perceptual Learning Reduces Interneuronal Correlations in Macaque Visual Cortex. Neuron, 71(4), 750–761. 10.1016/j.neuron.2011.06.015

International Brain Laboratory. (2020). Behavior: Appendix 2: IBL protocol for mice training. figshare. 10.6084/m9.figshare.11634729.v3

International Brain Laboratory, Aguillon-Rodriguez, V., Angelaki, D., Bayer, H., Bonacchi, N., Carandini, M., Cazettes, F., Chapuis, G., Churchland, A. K., Dan, Y., Dewitt, E., Faulkner, M., Forrest, H., Haetzel, L., Häusser, M., Hofer, S. B., Hu, F., Khanal, A., Krasniak, C., … Zador, A. M. (2021). Standardized and reproducible measurement of decision-making in mice. eLife, 10, e63711. 10.7554/eLife.63711

International Brain Laboratory, Banga, K., Benson, J., Bhagat, J., Biderman, D., Birman, D., Bonacchi, N., Bruijns, S. A., Buchanan, K., Campbell, R. A. A., Carandini, M., Chapuis, G. A., Churchland, A. K., Davatolhagh, M. F., Lee, H. D., Faulkner, M., Gerçek, B., Hu, F., Huntenburg, J., … Zhang, Y. (2025). Reproducibility of in vivo electrophysiological measurements in mice. eLife, 13, RP100840. 10.7554/eLife.100840

International Brain Laboratory, Meshulam, L., Angelaki, D., Benson, B., Benson, J., Birman, D., Bruijns, S. A., Carandini, M., Catarino, J. A., Chapuis, G. A., Churchland, A. K., Dan, Y., Davatolhagh, F., Dayan, P., DeWitt, E. E., Engel, T. A., Fabbri, M., Faulkner, M., Fiete, I. R., … Witten, I. B. (2025). A brain-wide map of neural activity during complex behaviour. Nature, 645(8079), 177–191. 10.1038/s41586-025-09235-0

Iurilli, G., & Datta, S. R. (2017). Population Coding in an Innately Relevant Olfactory Area. Neuron, 93(5), 1180–1197.e7. 10.1016/j.neuron.2017.02.010

Jun, J. J., Steinmetz, N. A., Siegle, J. H., Denman, D. J., Bauza, M., Barbarits, B., Lee, A. K., Anastassiou, C. A., Andrei, A., Aydın, Ç., Barbic, M., Blanche, T. J., Bonin, V., Couto, J., Dutta, B., Gratiy, S. L., Gutnisky, D. A., Häusser, M., Karsh, B., … Harris, T. D. (2017). Fully integrated silicon probes for high-density recording of neural activity. Nature, 551(7679), Article 7679. 10.1038/nature24636

Kim, Y., Yang, G. R., Pradhan, K., Venkataraju, K. U., Bota, M., Molino, L. C. G. del, Fitzgerald, G., Ram, K., He, M., Levine, J. M., Mitra, P., Huang, Z. J., Wang, X.-J., & Osten, P. (2017). Brain-wide Maps Reveal Stereotyped Cell-Type-Based Cortical Architecture and Subcortical Sexual Dimorphism. Cell, 171(2), 456–469.e22. 10.1016/j.cell.2017.09.020

Li, S.-C., & Rieckmann, A. (2014). Neuromodulation and aging: Implications of aging neuronal gain control on cognition. Current Opinion in Neurobiology, SI: Neuromodulation, 29, 148–158. 10.1016/j.conb.2014.07.009

Licata, A. M., Kaufman, M. T., Raposo, D., Ryan, M. B., Sheppard, J. P., & Churchland, A. K. (2017). Posterior Parietal Cortex Guides Visual Decisions in Rats. Journal of Neuroscience, 37(19), 4954–4966. 10.1523/JNEUROSCI.0105-17.2017

Liu, X., Zhou, Y., Liu, J., & Xu, G. (2025). Metformin improves age-related visual cortex dysfunction in mice by reducing noise correlation in the primary visual cortex. Frontiers in Aging Neuroscience, 17. 10.3389/fnagi.2025.1572653

MacDonald, S. W. S., Nyberg, L., & Bäckman, L. (2006). Intra-individual variability in behavior: Links to brain structure, neurotransmission and neuronal activity. Trends in Neurosciences, 29(8), 474–480. 10.1016/j.tins.2006.06.011

McGinley, M. J., Vinck, M., Reimer, J., Batista-Brito, R., Zagha, E., Cadwell, C. R., Tolias, A. S., Cardin, J. A., & McCormick, D. A. (2015). Waking State: Rapid Variations Modulate Neural and Behavioral Responses. Neuron, 87(6), 1143–1161. 10.1016/j.neuron.2015.09.012

Minces, V., Pinto, L., Dan, Y., & Chiba, A. A. (2017). Cholinergic shaping of neural correlations. Proceedings of the National Academy of Sciences, 114(22), 5725–5730. 10.1073/pnas.1621493114

Miura, K., Mainen, Z. F., & Uchida, N. (2012). Odor Representations in Olfactory Cortex: Distributed Rate Coding and Decorrelated Population Activity. Neuron, 74(6), 1087–1098. 10.1016/j.neuron.2012.04.021

Montijn, J. S., Vinck, M., & Pennartz, C. M. A. (2014). Population coding in mouse visual cortex: Response reliability and dissociability of stimulus tuning and noise correlation. Frontiers in Computational Neuroscience, 8. 10.3389/fncom.2014.00058

Moreno-Bote, R., Beck, J., Kanitscheider, I., Pitkow, X., Latham, P., & Pouget, A. (2014). Information-limiting correlations. Nature Neuroscience, 17(10), 1410–1417. 10.1038/nn.3807

Ni, A. M., Bowes, B. S., Ruff, D. A., & Cohen, M. R. (2022). Methylphenidate as a causal test of translational and basic neural coding hypotheses. Proceedings of the National Academy of Sciences, 119(17), e2120529119. 10.1073/pnas.2120529119

Ni, A. M., Ruff, D. A., Alberts, J. J., Symmonds, J., & Cohen, M. R. (2018). Learning and attention reveal a general relationship between population activity and behavior. Science, 359(6374), 463–465. 10.1126/science.aao0284

Poland, E., Donner, T. H., Müller, K.-M., Leopold, D. A., & Wilke, M. (2019). Thalamus exhibits less sensory variability quenching than cortex. Scientific Reports, 9(1), Article 1. 10.1038/s41598-019-43934-9

Ponce-Alvarez, A., Thiele, A., Albright, T. D., Stoner, G. R., & Deco, G. (2013). Stimulus-dependent variability and noise correlations in cortical MT neurons. Proceedings of the National Academy of Sciences, 110(32), 13162–13167. 10.1073/pnas.1300098110

Rabinowitz, N. C., Goris, R. L., Cohen, M., & Simoncelli, E. P. (2015). Attention stabilizes the shared gain of V4 populations. eLife, 4, e08998. 10.7554/eLife.08998

Ruff, D. A., & Cohen, M. R. (2014). Attention can increase or decrease spike count correlations between pairs of neurons depending on their role in a task. Nature Neuroscience, 17(11), 1591–1597. 10.1038/nn.3835

Ruff, D. A., & Cohen, M. R. (2016). Stimulus Dependence of Correlated Variability across Cortical Areas. Journal of Neuroscience, 36(28), 7546–7556. 10.1523/JNEUROSCI.0504-16.2016

Ryan, M. B., Ye, L., & Churchland, A. K. (2026). Harnessing theoretical neuroscience to understand decision-making across aging. Current Opinion in Neurobiology, 99, 103223. 10.1016/j.conb.2026.103223

Shi, Y.-L., Steinmetz, N. A., Moore, T., Boahen, K., & Engel, T. A. (2022). Cortical state dynamics and selective attention define the spatial pattern of correlated variability in neocortex. Nature Communications, 13(1), 44. 10.1038/s41467-021-27724-4

Shilling-Scrivo, K., Mittelstadt, J., & Kanold, P. O. (2021). Altered Response Dynamics and Increased Population Correlation to Tonal Stimuli Embedded in Noise in Aging Auditory Cortex. The Journal of Neuroscience, 41(46), 9650–9668. 10.1523/JNEUROSCI.0839-21.2021

Shilling-Scrivo, K., Mittelstadt, J., & Kanold, P. O. (2022). Decreased Modulation of Population Correlations in Auditory Cortex Is Associated with Decreased Auditory Detection Performance in Old Mice. Journal of Neuroscience, 42(49), 9278–9292. 10.1523/JNEUROSCI.0955-22.2022

Smith, M. A., & Kohn, A. (2008). Spatial and Temporal Scales of Neuronal Correlation in Primary Visual Cortex. The Journal of Neuroscience, 28(48), 12591–12603. 10.1523/JNEUROSCI.2929-08.2008

Smith, M. A., & Sommer, M. A. (2013). Spatial and Temporal Scales of Neuronal Correlation in Visual Area V4. Journal of Neuroscience, 33(12), 5422–5432. 10.1523/JNEUROSCI.4782-12.2013

Stringer, C., & Pachitariu, M. (2024). Analysis methods for large-scale neuronal recordings. Science, 386(6722), eadp7429. 10.1126/science.adp7429

Urai, A. E. (2026). Structure uncovered: Understanding temporal variability in perceptual decision-making. Trends in Cognitive Sciences, 30(1), 54–65. 10.1016/j.tics.2025.06.003

Urai, A. E., Doiron, B., Leifer, A. M., & Churchland, A. K. (2022). Large-scale neural recordings call for new insights to link brain and behavior. Nature Neuroscience, 25(1), Article 1. 10.1038/s41593-021-00980-9

Vinck, M., Batista-Brito, R., Knoblich, U., & Cardin, J. A. (2015). Arousal and locomotion make distinct contributions to cortical activity patterns and visual encoding. Neuron, 86(3), 740–754. 10.1016/j.neuron.2015.03.028

Wang, X., Zhang, B., Wang, H., Liu, J., Xu, G., & Zhou, Y. (2019). Aging affects correlation within the V1 neuronal population in rhesus monkeys. Neurobiology of Aging, 76, 1–8. 10.1016/j.neurobiolaging.2018.11.025

Welford, A. T. (1981). Signal, Noise, Performance, and Age. Human Factors: The Journal of the Human Factors and Ergonomics Society, 23(1), 97–109. 10.1177/001872088102300109

Xu, G., Liu, X., Liu, J., & Zhou, Y. (2026). GABAergic interneuron subtypes differentially regulate noise correlations that are critical for visual orientation discrimination in mice. Cell Reports, 45(1). 10.1016/j.celrep.2025.116805

Xue, B., Meng, X., Kao, J. P. Y., & Kanold, P. O. (2023). Age-related changes in excitatory and inhibitory intra-cortical circuits in auditory cortex of C57Bl/6 mice. Hearing Research, 429, 108685. 10.1016/j.heares.2022.108685

Zang, F., Khanal, A., Förster, S., Churchland, A. K., & Urai, A. E. (2026). Age-related changes in behavioural and neural variability in a decision-making task. Nature Communications, 17(1), 8156. 10.1038/s41467-026-74227-1

Zohary, E., Shadlen, M. N., & Newsome, W. T. (1994). Correlated neuronal discharge rate and its implications for psychophysical performance. Nature, 370(6485), 140–143. 10.1038/370140a0

